# Influence of Fara-darmani Consciousness Field on Bacterial Population Growth

**DOI:** 10.1101/2021.01.08.426007

**Authors:** Mohammad Ali Taheri, Gholamreza Zarrini, Sara Torabi, Noushin Nabavi, Farid Semsarha

## Abstract

The treatment of bacterial infections and the rising challenges of antibiotics resistance are global concerns and the primary topics in basic science and clinical microbiology. In the present study, the effectiveness of treatment of selected populations of bacteria using an immaterial and non-energetic method called Fara-darmani Consciousness Field treatment is investigated. Population growth was assessed by turbidimetry, colony counting and tetrazolium chloride reduction assays in non-treated control and Fara-darmani-treated groups. Our results suggest effectiveness of the Fara-darmani Consciousness Field on reducing various types of bacterial strain growth rates (up to 46%). In addition, along with a decrease in bacterial population, evidence of increased survival can be seen in the larger healthy population (up to about 60%). Thus, in this study, we confirm the effects of the Consciousness Field on bacterial population survival. This study also warrants additional research.

## 1. Introduction

Bacteria have existed before humans as the first forms of life on earth with the need to adapt to environmental conditions and changes in time. Many bacteria use ‘quorum sensing’ (QS) to control their gene expression in response to their population density. During specific stages of growth or other inputs like environmental stresses, the bacteria produce signaling molecules at a threshold concentration, which signal other regulators that can induce or repress target genes (Frederix and Dowine, 2011). Bacteria with development of communication capabilities (e.g. quorum sensing, chemotactic signaling and plasmid exchange), afford better adaptability to growth conditions (Ben-Jacob, 2003). Additionally, bacteria can form structured colonies to increase their benefits from accessing resources, a characteristic that individual bacterial cells cannot effectively utilize (Shapiro, 1998).

Much research has been performed to study bacterial growth properties and characteristics (Shaechter, 2015). It has been demonstrated that bacteria will growth in a predictable pattern, with four distinct phases, when placed in a suitable medium. Initially, the bacteria will grow rather slowly (lag phase), before reaching a maximum growth rate with greater rapidity (log phase). Following that, bacteria reach a plateau phase where the rate of growth and death become equal (stationery phase). In the final decline phase, the rate of cell death exceeds the rate of growth. The growth curve of bacterial population is similar to other living populations in a restricted area (Henrici, 1928). In a way, bacteria have the ability of ‘linguistic’ communication and social intelligence (Jacob et al, 2004).

Antibiotics have been developed to combat disease-causing bacteria such as infections, tuberculosis, gonorrhea, plague, or anthrax among others. However, bacteria have the ability to become resistant to antibiotics under prolonged selection pressures. According to the Center for Disease Control and Prevention (CDC), antibiotic resistance is one of the most serious health treats. Each year in the U.S., at least 2.8 million people get an antibiotic-resistance infection, and more than 35,000 people die. Due to the increase in resistance rates to conventional antibiotics, alternatives methods such as bacteriophage therapy (Golkar et al, 2014), predatory bacteria (Kadouri et al, 2013), or bacteriocins (Cotter et al, 2013) are being investigated. Additionally, many varieties of compounds produced by plants have proved to have therapeutic potentials and antimicrobial effects or elicit modifications to antibiotic resistance (Sibanda and Okoh, 2007). Another alternative for growth inhibition of resistant bacteria is attenuation of bacterial virulence by inactivating the QS system of a pathogen (Hentzer and Givskov, 2003).

Very little information is available on the complementary therapy methods that can induce changes in bacterial population growth status in culture media. The Fara-darmani Consciousness Field (CF) as a complementary therapy was founded by Mohammad Ali Taheri. In this theoretical concept, cosmic consciousness is the collection of consciousness, wisdom or intelligence governing the world of existence. According to Taheri (2013), there are variable CFs and Fara-darmani CF is a subcategory of the Cosmic Consciousness Network (CCN). Consciousness is one of the three existing elements of the universe different from matter and energy. Assuming consciousness as neither matter nor energy, we cannot measure it quantitatively and can only screen its effects indirectly through experimentation. In this experimentation, mind-matter interaction occurs by connecting to CCN. A Fara-therapist’s mind establishes a consciousness bond between the whole consciousness (CCN) and the subjects of study through which all constituents will get scanned and corrected. In this theory, any living creature including humans, plants, animals, or microorganisms, can be cured via humans by connecting to CCN through a Fara-therapist’s mind (Taheri, 2013). In previous studies, the effects of Fara-darmani CF was investigated on change in cancer cell growth (Taheri et al, 2020^a^), electrical activity in the brain of Fara-therapists (Taheri et al 2020^b^) and wheat plants (Torabi et al, 2020). In this study, the influence of Fara-darmani Consciousness Field on the growth behavior of different populations of laboratory and nosocomial bacterial strains are investigated.

## 2. Materials and Methods

### 2.1 Fara-darmani Consciousness Field application

The use of Fara-darmani Consciousness Field is done according to the protocols mentioned in the website of research management in the Consciousness Field according to Taheri (www.cosmointel.com). After visiting the website and requesting the application of Consciousness Field treatment for the subject of study (in the assign announcement section), according to the time and place determined by the researcher of the study, the treatments are allocated to the study for free. The main requirement of this method, according to the principles introduced by the founder of this method, is to establish a relationship between the part (subject of study) and whole consciousness (CCN) through the human mind (Fara-therapist’s mind). After communicating with CF according to the time and place specified by the researcher, and assigning the treatment to the study, the possible results can be analyzed and evaluated in accordance with the consequent methodologies.

### 2.2 Turbidimetry of the primary selected bacterial strains at 24 hours

Bacterial growth under the influence of Fara-darmani CF were measured by turbidimetry at OD600 nm in tube cultures. For this purpose, 36 test tubes containing 10 ml of Müller-Hinton Broth medium were prepared and autoclaved. Two sets of the culture tubes inoculated with 10^2^ and 10^5^ cfu/ml of test microorganisms. Control and treatment samples were placed in a separate holder in an incubator on different levels. Sampling of cultures was done at 24 hours after the start of incubation with a volume of 1.5 ml.

### 2.3 Bacterial growth analysis at different time intervals

For additional interpretation of the growth behavior of bacteria under treatment, bacterial growth changes were measured by three methods: (1) turbidimetry at OD 600 nm, (2) colony counting, and (3) assay of the bacterial regenerative power in reduction of tetrazolium chloride. To evaluate the effect of Fara-darmani CF in different time intervals, sampling was done three times in two different experiment steps: in first step, at 6, 16 and 24 hours and in the second step at 1, 3 and 6 hours.

In this experiment section and in the first step, 8 test tubes containing 10 ml of Müller-Hinton Broth medium and 8 Erlenmeyer 100 containing 20 ml of Müller-Hinton Broth were prepared and autoclaved. For each of the studied microorganisms in step 1, one tube and one Erlenmeyer flask were considered as the control group and one tube and Erlenmeyer flask was considered as the treatment group. In step 2, only growth in the Erlenmeyer flask was considered.

One ml of bacteria with 10^5^ cfu/ml was added to each tube and Erlenmeyer flask culture medium. Control and treatment samples were placed in a separate holder in an incubator on the different levels. Erlenmeyer flasks were placed in shaker incubators with separate rows and the longest distance between control and treatment samples. Sampling of cultures was done at 6, 16 and 24 hours (in step 1) and at 1, 3 and 6 hours (in step 2) after the start of incubation with a volume of 1.5 ml. Turbidity was measured at 600 nm (for step 1 of this experiment section) and surface culture was done for colony count (in two replications) for each culture medium in the two mentioned steps. For cell survival assay by tetrazolium chloride, in step 1, we added 10 µl of 1 mg/ml aqueous solution to 1 ml of microbial sample and after one hour of incubation at 35 °C, the absorbance of the samples was read at 495 nm. In step 2, all procedures were similar to step 1, except that the concentration of tetrazolium chloride was used at 100 times more than step 1.

## Results

### 3.1 CF Effect on the primary selected bacterial strains at 24 hours

In order to investigate the effect of Fara-darmani CF on bacteria, four laboratory strains and five nosocomial strains were used. Efficacy was reported based on the percentage of reduction of microbial populations as shown in Table 1.

**Table 1.** Absorption change at 600 nm for tube bacterial culture at the 24 hour.

| Type of strain | Strain* | Characteristics | %Reduction<br>(Early population: 10<sup>2</sup>) | %Reduction<br>(Early population: 10<sup>5</sup>) |
| --- | --- | --- | --- | --- |
| Laboratory | *E.coli* | Symbiotic | 18.38 | 8.7 |
|  | *K.pneumoniae*(1) | Symbiotic | 33.78 | 17.32 |
|  | *S.aureus*(1) | Symbiotic | 24.17 | 45.04 |
|  | *B.subtillis* | Environmental | 24.4 | 26.98 |
| Nosocomial | *P.aeruginosia*(1) | Pathogenic | 4.2 | 6.2 |
|  | *P.aeruginosia*(2) | Pathogenic | 1.7 | 12.7 |
|  | *K.pneumoniae*(2) | Pathogenic | 5.6 | 0.7 |
|  | *S.aureus*(2) | Pathogenic | 5.4 | 0.6 |
|  | *S.aureus*(3) | Pathogenic | 7.6 | 1 |

\* The numbers in parentheses refer to different strains of a bacterial species.

According to the results presented in Table 1, the highest decrease in the percentage of laboratory microbial populations was observed in the 10^5^ cfu/ml concentration of *S*.*aureus* and the lowest decrease was observed in the same concentration in the case of *E*.*coli*. The varying initial microbial populations of gram-positive or negative bacteria do not show a significant difference in the results. Moreover, in nosocomial bacteria, the highest decrease in the percentage of microbial population was observed in the 10^5^ cfu/ml concentration of *P*.*aeruginosa (2)* and the lowest decrease in the same concentration in the case of *K*.*pneumonia*. Overall, according to the results, the effectiveness of the CF on nosocomial strains is less than that of laboratory strains.

### 3.2 CF influence on bacterial growth at different time intervals

Four bacteria, *Escherichia coli, Pseudomonas aeruginosa, Staphylococcus aureus* and *Bacillus subtilis* were used to study the effect of CF treatment on the growth behavior of bacteria. In the first step, growth of the selected strains was examined at 6, 16 and 24 hours in both tube and Erlenmeyer flask culture of bacteria through turbidity measurements, colony counting and tetrazolium chloride reduction methods. After consideration of the results of the first step of this section, in the second step, similar studies were done at 1, 3 and 6 hours in Erlenmeyer flask condition through colony counting and tetrazolium chloride method as discussed below.

#### 3.2.1 Turbidity measurements

In order to investigate the results of section 3.1 and to study changes in populations at different times, we assessed bacterial growth by the turbidimetric method at 6h, 16h and 24h. The tube method is the conventional way for studying the antimicrobial effects of antibiotics, whereas the Erlenmeyer flask provides a more suitable growth medium for bacteria due to better aeration and uniformity of the growth medium. In this study, to determine the effect of treatment, both tube and Erlenmeyer methods were used, and the results are shown in tables 2 and 3.

**Table 2:** Absorption change at 600 nm for tube bacterial culture at 6, 16 and 24 hours.

| Type of strain | Strain | OD<sub>600nm</sub> |  |  |  |  |  | Efficacy of treatment (%) |  |  |
| --- | --- | --- | --- | --- | --- | --- | --- | --- | --- | --- |
|  |  | 6 hours |  | 16 hours |  | 24 hours |  | (-): Decrease in population<br>(+): Increase in population |  |  |
|  |  | Control | Treatment | Control | Treatment | Control | Treatment | 6h | 16h | 24h |
| Laboratory | *E. coli* | 0.168 | 0.121 | 0.436 | 0.409 | 0.878 | 0.950 | -27 | -6 | +8 |
|  | *B. subtilis* | 0.023 | 0.021 | 0.161 | 0.148 | 0.333 | 0.282 | -8 | -8 | -15 |
| Nosocomial | *P. aeruginosa* | 0.121 | 0.113 | 0.379 | 0.125 | 0.985 | 0.853 | -7 | -67 | -13 |
|  | *S. aureus* | 0.152 | 0.132 | 0.157 | 0.135 | 0.195 | 0.192 | -13 | -14 | -1.5 |

**Table 3:**
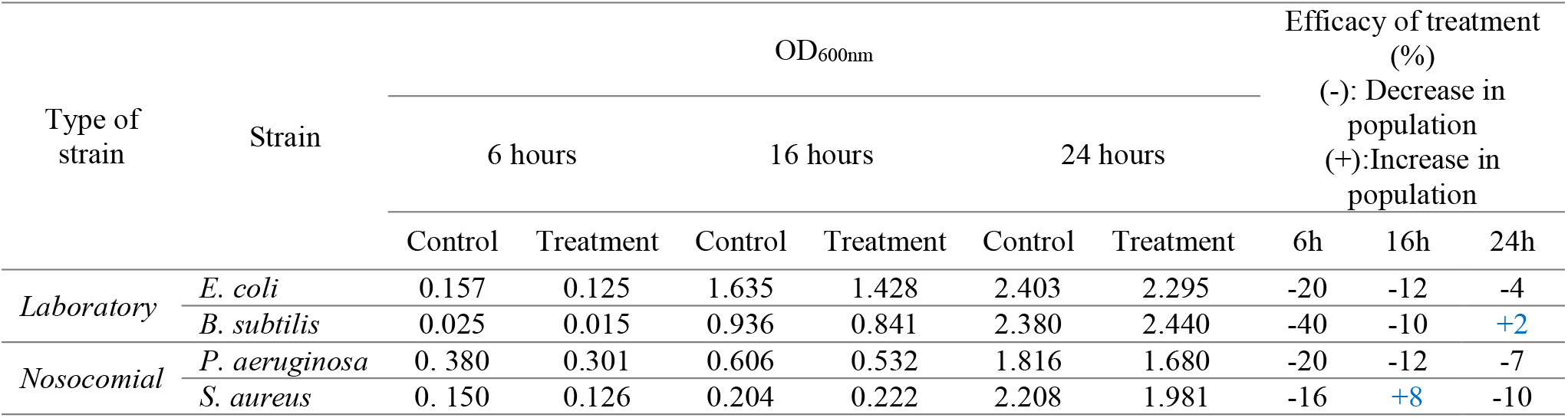
Absorption change at 600 nm for bacterial culture in Erlenmeyer flask culture at 6, 16 and 24 hours.

| Type of strain | Strain | OD<sub>600nm</sub> |  |  |  |  |  | Efficacy of treatment (%) |  |  |
| --- | --- | --- | --- | --- | --- | --- | --- | --- | --- | --- |
|  |  | 6 hours |  | 16 hours |  | 24 hours |  | (-): Decrease in population<br>(+): Increase in population |  |  |
|  |  | Control | Treatment | Control | Treatment | Control | Treatment | 6h | 16h | 24h |
| Laboratory | *E. coli* | 0.157 | 0.125 | 1.635 | 1.428 | 2.403 | 2.295 | -20 | -12 | -4 |
|  | *B. subtilis* | 0.025 | 0.015 | 0.936 | 0.841 | 2.380 | 2.440 | -40 | -10 | +2 |
| Nosocomial | *P. aeruginosa* | 0.380 | 0.301 | 0.606 | 0.532 | 1.816 | 1.680 | -20 | -12 | -7 |
|  | *S. aureus* | 0.150 | 0.126 | 0.204 | 0.222 | 2.208 | 1.981 | -16 | +8 | -10 |

The results of tube and Erlenmeyer flask culture by turbidity measurement show that there are differences in the absorption of control and treatment samples. The difference in absorption for the 6-hour culture time is greater than the 16-hour and 24-hour culture time. The difference in absorption in 6 hours after culture shows a greater difference, which can be due to the greater impact of CF treatment on bacteria during this time. This trend can also be compensated by the growth of bacteria in the continuation of cultivation. Interestingly, we observed an increase in growth and colony counts (number of colonies) of the reduced population of bacteria in cases such as the 24-hour culture of *E*.*coli* and *B*.*subtilis*, in both tube and Erlenmeyer flask cultures.

#### 3.2.2 Colony count measurements

Since live and dead bacteria are indistinguishable in the turbidity method, colony counting method was used to measure the status of bacteria in the culture medium. For this purpose, in the first step, tube and Erlenmeyer flask cultures were cultured at 6, 16 and 24 hours in Müller-Hinton medium with two replications and then counted. The results of colony count are given in Tables 4 and 5.

**Table 4:** Colony count results of treatment and control samples in tube culture method at 6, 16 and 24 hours.

| Type of strain | Strain | Colony Count |  |  |  |  |  | Efficacy of treatment (%) |  |  |
| --- | --- | --- | --- | --- | --- | --- | --- | --- | --- | --- |
|  |  | 6 hours/10<sup>4</sup> |  | 16 hours/ 10<sup>6</sup> |  | 24 hours |  | (-): Decrease in population<br>(+): Increase in population |  |  |
|  |  | Control | Treatment | Control | Treatment | Control | Treatment | 6h | 16h | 24h |
| Laboratory | *E. coli* | NC* | NC | 357 | 285 | NC | NC | ND** | -20 | ND |
|  | *B. subtilis* | 55 | 40 | 20 | 16 | NC | NC | -27 | -20 | ND |
| Nosocomial | *P. aeruginosa* | 398 | NC | 49 | 41 | NC | NC | ND | -16 | ND |
|  | *S. aureus* | 310 | 184 | 37 | 29 | NC | NC | -39 | -25 | ND |

NC \* means uncountable; ND\*\* Not determined

**Table 5:** Colony count results of treatment and control samples in Erlenmeyer flasks at 6, 16 and 24 hours.

| Type of strain | Strain | Colony Count |  |  |  |  |  | Efficacy of treatment (%) |  |  |
| --- | --- | --- | --- | --- | --- | --- | --- | --- | --- | --- |
|  |  | 6 hours/10<sup>4</sup> |  | 16 hours/10<sup>6</sup> |  | 24 hours/10<sup>8</sup> |  | (-): Decrease in population<br>(+): Increase in population |  |  |
|  |  | Control | Treatment | Control | Treatment | Control | Treatment | 6h | 16h | 24h |
| Laboratory | *E. coli* | NC | 1328 | 433 | 320 | NC | NC | ND | -26 | ND |
|  | *B. subtilis* | 190 | 124 | 51 | 45 | NC | NC | -34 | -11 | ND |
| Nosocomial | *P. aeruginosa* | 169 | 179 | 487 | 352 | NC | NC | +6 | -27 | ND |
|  | *S. aureus* | NC | 100 | 64 | 53 | NC | NC | ND | -16 | ND |

Colony count results for 24-hour samples was not possible due to overgrowth (NC) and in cases where the count numbers of one of the control or treatment samples were not obtained, it was not possible to determine the percentage of treatment efficiency. In other cases, the results of colony counts showed the effectiveness of the Fara-darmani CF treatment on declining the bacterial population, although in the case of *P. aeruginosa* at 6 hours, the opposite result (increase in population) was observed.

In step 2 and after observing the effects of the Fara-darmani Consciousness Field in the first step in turbidity measurement and colony counting, the colony counting of Erlenmeyer flask samples were repeated and sampling was done at 1, 3 and 6 hours. The results of this step are given in Table 6.

**Table 6.** Colony counting of Fara-darmani CF treatment and control samples in Erlenmeyer flasks at 1, 3 and 6 hours.

| Type of strain | Strain | Colony Count |  |  |  |  |  | Efficacy of treatment (%) |  |  |
| --- | --- | --- | --- | --- | --- | --- | --- | --- | --- | --- |
| | | 1 hours/$10^4$ | | 3 hours/$10^4$ | | 6 hours/$10^5$ | | (-): Decrease in population<br>(+): Increase in population | | |
|  |  | Control | Treatment | Control | Treatment | Control | Treatment | 1h | 3h | 6h |
| Laboratory | *E. coli* | 14 | 9 | 105 | 78 | 538 | 397 | -35 | -23 | -26 |
|  | *B. subtilis* | 3.4 | 2.2 | 6.2 | 4.3 | 9.6 | 7.5 | -34 | -30 | -22 |
| Nosocomial | *P. aeruginosa* | 26 | 13.9 | 77 | 48 | 293 | 222 | -46 | -37 | -24 |
|  | *S. aureus* | 11.9 | 9.1 | 35 | 24 | 88 | 73 | -23 | -31 | -17 |

As can be seen in Table 6, the effectiveness of the Fara-darmani CF treatment in the first hour of study is greatest in the two laboratory-strains and in *P. aeruginosa*. Moreover, *S. aureus* shows the greatest reduction up to 3 hours. These results are in agreement with Table 2 which shows a decrease even in the early treatment times.

As shown in figure 1, the exponential growth plot in the first 6 hours of the Fara-darmani CF treatment and control population shows a significant and visible effect of this field on the various bacterial populations in the first 1 to 3 hours of the bacterial growth.

**Figure 1.**
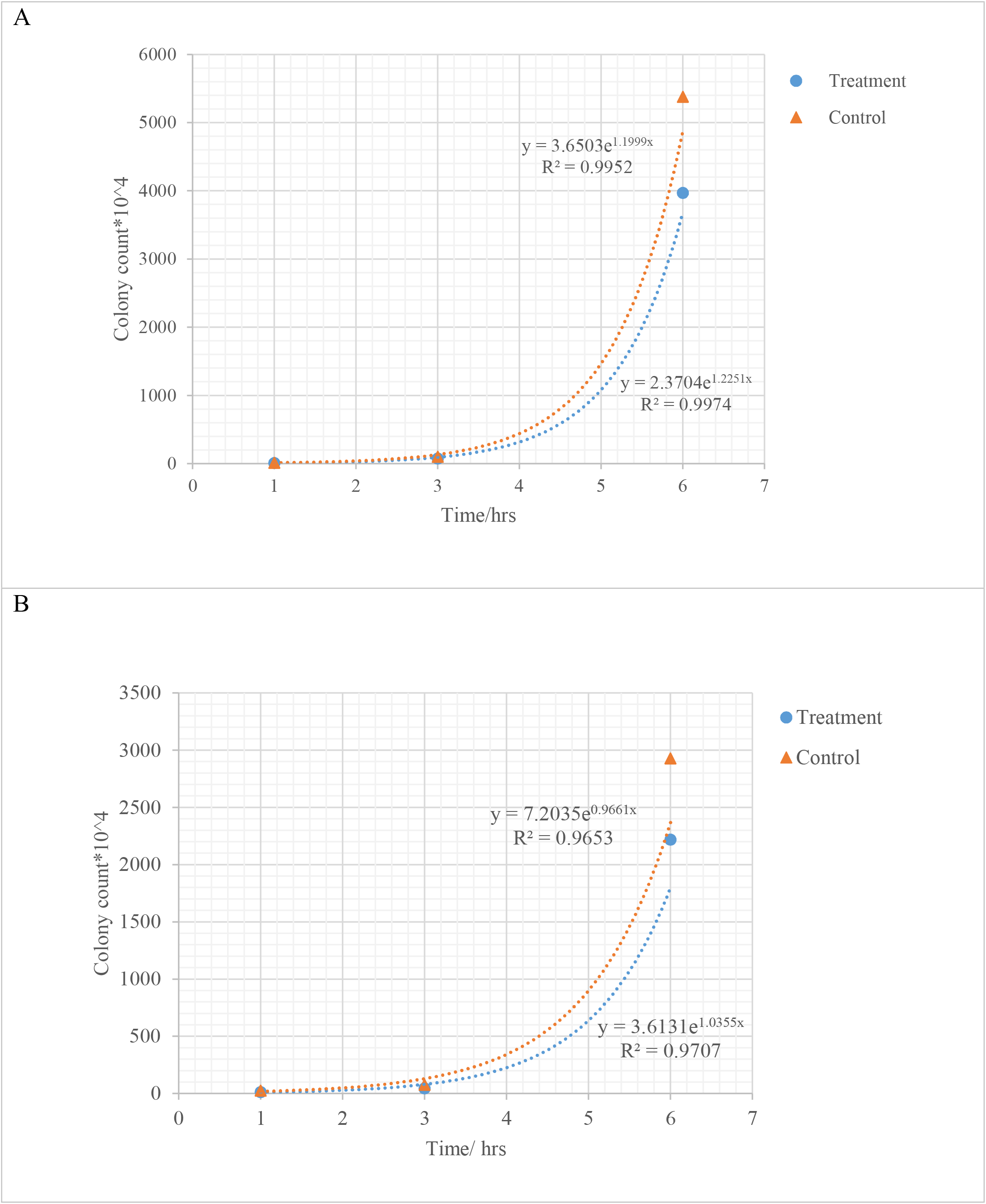
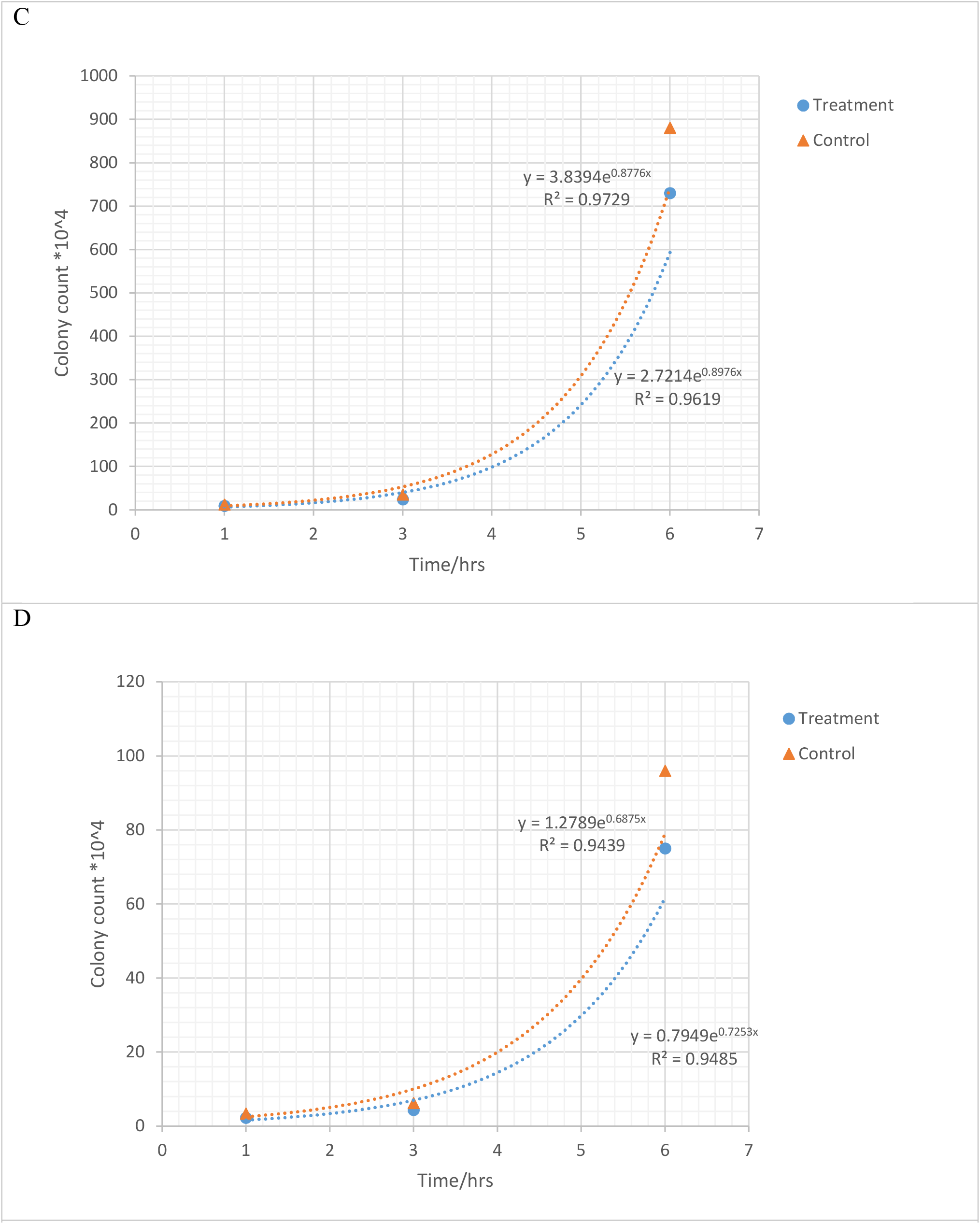
Exponential growth plot in the first three hours of the bacterial life cycle of CF treatmnet sample in comparison with the control samples.C: *P. aeruginosa*. D: *S. aureus*.

#### 3.2.3 Tetrazolium chloride reduction assay

Tetrazolium chloride compound was used to evaluate the metabolic status of bacteria and possible changes in bacterial regenerative power that indicate cell viability. Reduction of tetrazolium chloride by dehydrogenase enzymes in healthy bacterial cells produces a red color that is capable of absorbing light at 495 nm. For the tetrazolium chloride method in the first step, in 6 and 16 hours, we did not record a measurement due to low microbial concentration and low color. The earliest time a measurement was possible was at the 24 hours as shown in Table 7.

**Table 7:** Tetrazolium chloride reduction assay of tube and Erlenmeyer flask bacterial culture by absorbing light at 495 nm at the 24 hour.

| Type of strain | Strain | Tube culture |  |  | Erlenmeyer flask culture |  |  |
| --- | --- | --- | --- | --- | --- | --- | --- |
|  |  | A<sub>495nm</sub> |  | Efficacy of treatment (%)<sup>*</sup> | A<sub>495nm</sub> |  | Efficacy of treatment (%)<sup>*</sup> |
|  |  | Control | Treatment |  | Control | Treatment |  |
| *Laboratory* | *E. coli* | 1.7519 | 1.1173 | -36 | 1.1426 | 1.0929 | -4 |
|  | *B. subtilis* | 0.4960 | 0.7915 | +59 | 0.3839 | 0.2790 | -27 |
| *Nosocomial* | *P. aeruginosa* | 0.6679 | 0.7043 | +5 | 2.4616 | 2.0637 | -16 |
|  | *S. aureus* | 0.6063 | 0.7886 | +30 | 0.7366 | 0.6544 | -11 |

\* +: Increase in population; -: Decrease in population

The Erlenmeyer flask culture shows a more consistent result with the population decline in the previous methods. However, the results of this method for tube culture of bacteria, are consistent with previous results, and show a higher growth rate in the treated microbial samples than in the control (compared to efficacy results in the tables 2, 3 and 5). Tetrazolium chloride assay was also done on bacterial populations at 1, 3 and 6 hours as shown in Table 8.

**Table 8.** Tetrazolium chloride reduction assay of Erlenmeyer flask bacterial culture by absorbing light at 495 nm at 1, 3 and 6 hours.

| Type of bacteria | Strain | A<sub>495nm</sub> |  |  |  |  |  | Efficacy of treatment (%) |  |  |
| --- | --- | --- | --- | --- | --- | --- | --- | --- | --- | --- |
|  |  | 1 hours |  | 3 hours |  | 6 hours |  | 1h | 3h | 6h |
|  |  | Control | Treatment | Control | Treatment | Control | Treatment |  |  |  |
| *Laboratory* | *E. coli* | 0.044 | 0.032 | 0.094 | 0.083 | 0.633 | 0.534 | -27 | -11 | -15 |
|  | *B. subtilis* | 0.039 | 0.029 | 0.078 | 0.054 | 0.095 | 0.079 | -25 | -30 | -17 |
| *Nosocomial* | *P. aeruginosa* | 0.058 | 0.046 | 0.085 | 0.076 | 0.169 | 0.134 | -20 | -10 | -20 |
|  | *S. aureus* | 0.054 | 0.037 | 0.067 | 0.063 | 0.094 | 0.086 | -31 | -5 | -8 |

In Table 8, the longer time in the Tetrazolium chloride reduction assay causes the greatest decrease in populations and survival occurs only in the first 1 hour of the experiment (in the case of *B. subtilis* in the first 3 hours).

Figure 2 depicts exponential and logarithmic graphs of reduction rates of bacterial growth in tetrazolium chloride assay. Similar to the data obtained from other measurement methods for the same strains, we observe a general reduction in regenerative capacity in the Erlenmeyer culture treatment samples. The increase in regenerative capacity at 3 hours, in the case of *S. aureus* treatment, indicates better survival condition of the remaining bacterial populations, when compared to the tube culture of *B. subtilis, P. aeruginosa and S. aureus* strains.

**Figure 2.**
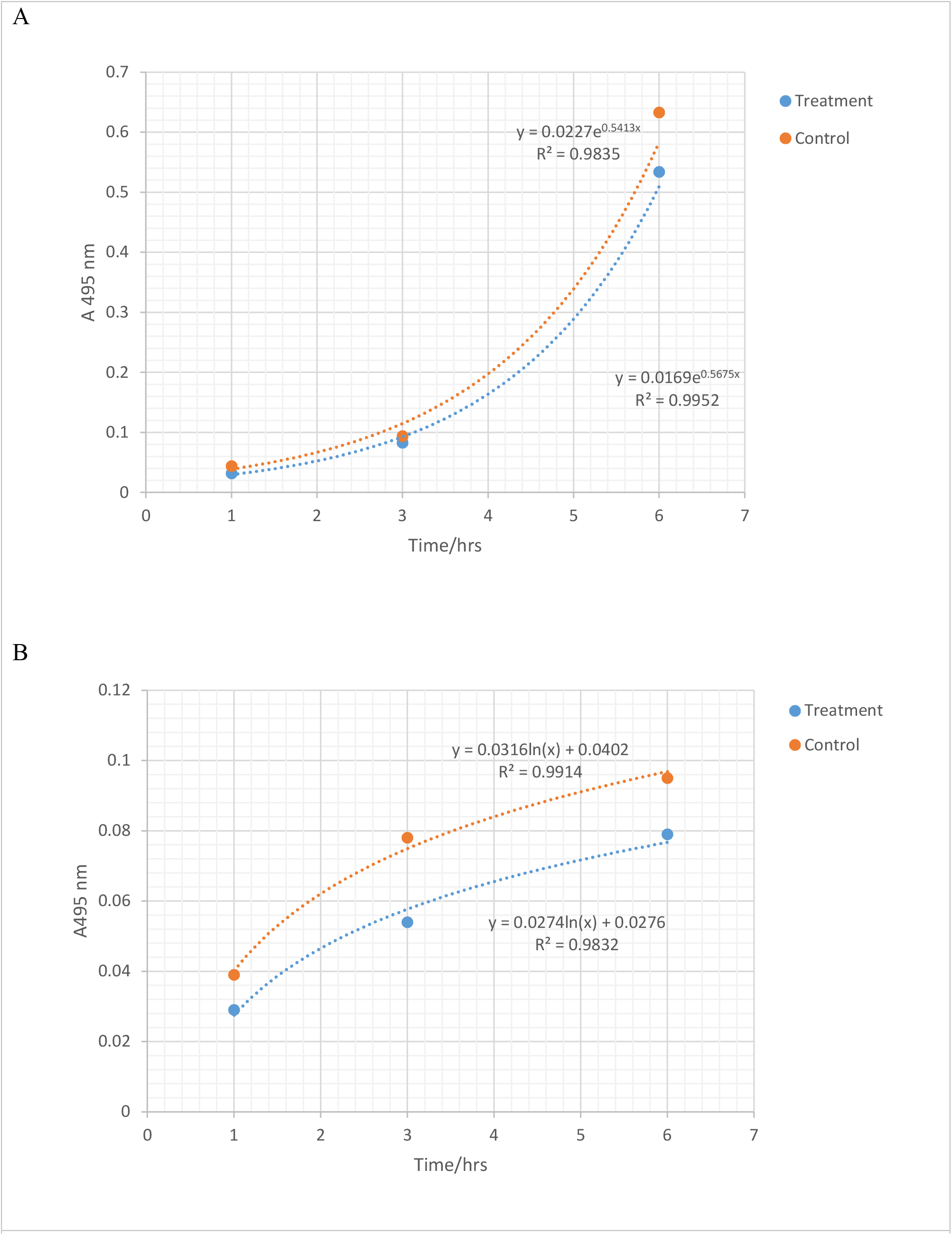
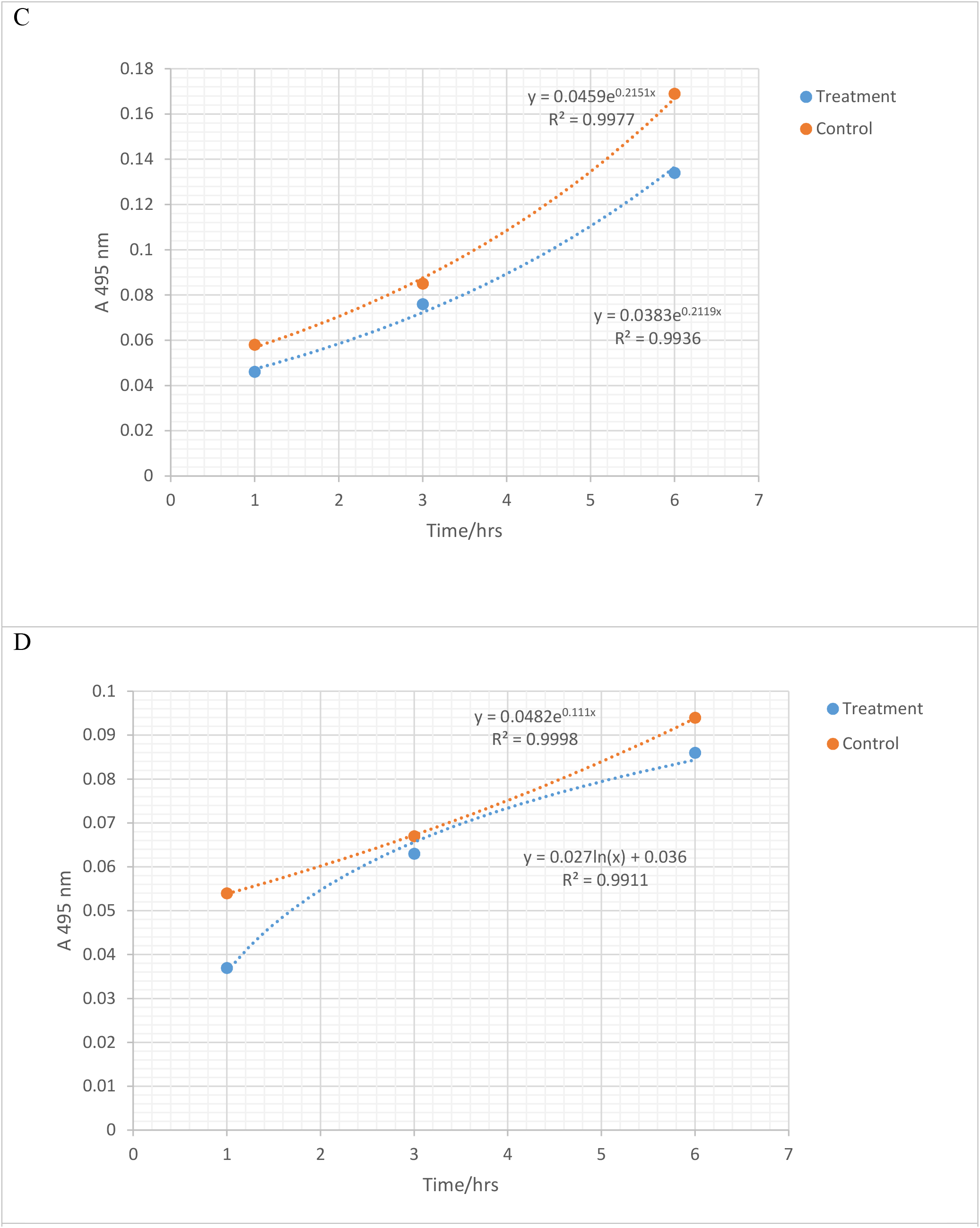
Change in thetrazolium choloride reduction in the first three hours of the bacterial life cycle of CF treated sample in comparison with the control samples. A:*E*.*coli*. B.*B*.*subtilis*. Change in thetrazolium choloride reduction in the first three hours of the bacterial life cycle of CF treated sample in comparison with the control samples. C: *P. aeruginosa*. D: *S. aureus*.

## 4. Discussion

Changes in the bacterial populations under the Fara-darmani Consciousness Field as mediated by the human mind is a novel study. The experiments in this study were initially performed on diverse bacterial populations to evaluate the initial effectiveness of the Consciousness Field 24 hours after the start of cultivation and treatment. In order to examine the reproducibility of the results and better interpret them, strains were selected from these tests, and at other intervals of bacterial growth cycle (6 to 24 hours) through sampling, repetition, and completion of growth tests. Finally, in the last stage, after confirming the observations in the previous stages, a supplementary study was performed to investigate the effects of the consciousness field, at shorter intervals of the bacterial life cycle (less than 6 hours).

According to Taheri, consciousness fields are immaterial and non-energetic fields with the ability to affect a variety of living and non-living systems from atom, to cells, to organisms. The general functions of these fields is to establish a connection between the subject under study and the Cosmic Consciousness Network, with the aim of reconstructing, modifying and repairing in order to achieve the optimal structure and performance of the system under study in its environment. What is observed in the present study is the reproducibility of a significant effect of the Consciousness Field on bacterial population growth. This effect occurs at first glance, with a decrease in the growth. In closer inspection, we found a concomitant change in the remained bacterial populations who have a higher ability to live and survive. In examining the results of the effect of Fara-darmani Consciousness Field on bacterial population growth and comparing it with respective control groups, the following summaries can be made: (1) The Fara-darmani Consciousness Field affects the selected population of bacteria in this study. This effect has been proved by studying different types of strains and repeating the study and sampling at different times and using complementary live and dead assay methods; (2) The effect of Consciousness Field treatment begins in the first hour of bacterial culture, simultaneously with the start of treatment and the synchronicity between the treatment and its effectiveness can be observed; (3) This effect has two manifestations: (a) population declines up to 46% in different bacteria (related to the origin and role in the ecosystem) and its evidence in both tube and Erlenmeyer culture media at different times of sampling (different stages of the bacterial life cycle), and (b) increase in the ability to regenerate and survive in the remaining bacterial populations, which in the conditions of tube culture are up to 60% in different bacterial strains; (4) laboratory bacterial strains show a greater decrease in growth than nosocomial strains with no significant difference in the initial population of bacteria nor in their gram-positive or negative characteristics; (5) Changes in environmental conditions (comparison of tube culture and Erlenmeyer culture) show the effectiveness of the Consciousness Field differently: the tube culture conditions, which are considered harsh environmental conditions for bacterial life, perform better than Erlenmeyer environment in showing the effect of the Consciousness Field on bacterial survival.

In order to further investigate bacterial populations affected by the consciousness fields, studies on other bacterial strains and especially on the antibiotic resistance of important nosocomial resistance bacterial strains is strongly recommended. Also, the use of other Consciousness Field types to observe the metabolic and physiological changes of bacteria seems to be fully justified.

